# PLANET: A Multi-Objective Graph Neural Network Model for Protein–Ligand Binding Affinity Prediction

**DOI:** 10.1101/2023.02.01.526585

**Authors:** Xiangying Zhang, Haotian Gao, Haojie Wang, Zhihang Chen, Zhe Zhang, Xinchong Chen, Yan Li, Yifei Qi, Renxiao Wang

**Author notes:** These authors had equal contributions to this work (Xiangying Zhang, Haotian Gao). To whom all correspondence should be addressed: (Y. Qi); (R. Wang).

## Abstract

Predicting protein-ligand binding affinity is a central issue in drug design. Various deep learning models have been developed in recent years to tackle this issue, but many of them merely focus on reproducing the binding affinity of known binders. In this study, we have developed a graph neural network model called PLANET (Protein-Ligand Affinity prediction NETwork). This model takes the graph-represented 3D structure of the binding pocket on the target protein and the 2D chemical structure of the ligand molecule as input, and it was trained through a multi-objective process with three related tasks, including deriving the protein–ligand binding affinity, protein–ligand contact map, and intra-ligand distance matrix. To serve those tasks, a large number of decoy non-binders were selected and added to the standard PDBbind data set. When tested on the CASF-2016 benchmark, PLANET exhibited a scoring power comparable to other deep learning models that rely on 3D protein–ligand complex structures as input. It also showed notably better performance in virtual screening trials on the DUD-E and LIT-PCBA benchmark. In particular, PLANET achieved comparable accuracy on LIT-PCBA as the conventional docking program Glide. However, it only took less than 1% of the computation time required by Glide to finish the same job because it did not perform exhaustive conformational sampling. In summary, PLANET exhibited a decent performance in binding affinity prediction as well as virtual screening, which makes it potentially useful for drug discovery in practice.

## 1. Introduction

Binding affinity to the chosen target protein has been an essential indicator in structure-based drug design for evaluating designed ligand molecules. Although protein-ligand binding affinity can be measured by various experimental assays, such assays are relatively expensive and often troublesome to set up. Besides, availability of the relevant protein and ligand samples is also an important issue. In contrast, computational methods are efficient, cost-effective, and do not need samples to work. Notably, the accuracy of free energy perturbation methods in binding affinity prediction gets close to the level of experimental errors in certain scenarios ^1–4^. However, application of such methods to high-throughput jobs, such as virtual screening, remains impractical. Combination of molecular docking and scoring function can achieve a balance between accuracy and efficiency, which is the common choice for jobs in reality ^5^. The accuracy of traditional scoring functions such as X-Score ^6^ and AutoDock Vina ^7^ is limited by the simple form of linear functions. Machine learning methods without predefined formulas can be more flexible to fit the training data and obtain higher accuracy than traditional scoring functions ^8^. Δ_Vina_RF_20_ ^9^, a random forest (RF)–based model, achieved the best metrics in almost all tests in the CASF-2016 benchmark ^10^. Machine learning models require feature engineering to transform the raw input to feature descriptors according to predefined rules, which need domain knowledge and can directly affect model performance, preventing end-to-end learning.

In contrast, deep learning models can make predictions based on the most related features automatically extracted from raw input and deep learning models have thus become a research hotspot in binding affinity prediction ^11, 12^. Some of those deep learning models do not rely on three-dimensional (3D) structures while the others do rely on 3D structures as inputs. Models in the first category ^13–19^ extract features from one-dimensional (1D) or two-dimensional (2D) structural inputs, such as protein sequences, SMILES strings, or 2D structural graphs. However, protein-ligand interactions occur in the 3D space, it is uncertain whether this level of information can be sufficiently degraded into lower dimensions and are thereby learned by this category of models. Models based on 3D structures directly take complex structures as inputs and one of the most common methods uses a 3D-convolutional neural network (CNN) ^20–29^ to extract features from binding sites that have been partitioned into small cubic grids, in which different channels encode structural information such as the positions of different types of atoms. Graph neural network (GNN)-based models ^30–33^ have also been proposed for predicting binding affinity. Tested on the CASF-2016 benchmark ^10^, most of these models can achieve higher scoring power than classical scoring functions. In addition to scoring power, screening power is also essential for binding affinity prediction models, that is, they should accurately rank binders and non-binders. The screening power determines whether these models can be used in virtual screening, which is one of the most important application scenarios for binding affinity prediction models. Unfortunately, the screening power of many models is less desirable or even being ignored ^34–36^, limiting their use in real-life drug design.

To improve the screening capacity of binding affinity prediction models, we have developed PLANET (i.e. Protein-Ligand Affinity prediction NETwork). PLANET is essentially a GNN model that takes the 3D structure of the binding pocket on the target protein and the 2D structural graph of the ligand molecule as inputs. It was trained through a multi-objective process as multi-objective training has been proven useful for improving the performance and generalization of binding affinity prediction models ^37^. The three basic objectives in our model training included deriving a protein– ligand contact map, intra-ligand distance matrix, and protein–ligand binding affinity. Deriving the protein–ligand contact map allows PLANET to capture protein–ligand interactions from the input structures, while deriving the intra-ligand distance matrix helps PLANET to capture 3D features from the 2D structural graph of the ligand. To reach these objectives, besides the commonly used PDBbind data set, artificially generated decoy molecules ^38^ were also added to the model training process. As tested on the CASF-2016 benchmark, PLANET exhibited a scoring power comparable to that of machine learning models that rely on 3D protein–ligand complex structures as inputs. As tested on the DUD-E ^39^ and LIT-PCBA ^40^ data sets, PLANET exhibited better screening power than other deep learning models trained on the PDBbind data set ^34^. Its performance in virtual screening trials was close to that of the popular convention docking method Glide, while its computational cost was only a tiny fraction of that of Glide.

## 2. Methods

### 2.1 Model Architecture

Our PLANET model has three major objectives:

#### (1) Prediction of protein–ligand binding affinity

PLANET relies on the 3D structure of the binding pocket on a given target protein and the 2D molecular graph i.e. chemical structure, of a given ligand molecule to predict the binding affinity between them. It makes prediction by integrating the contributions from pocket residues and ligand atoms that are involved in interaction. It is of course more straightforward to retrieve protein-ligand interactions from a 3D protein-ligand complex structure, just like the method employed by Zhu et al. in their work ^41^. However, the input used by PLANET does not provide such information directly. This difficulty requires PLANET to predict protein–ligand interactions and results into the second objective of PLANET.

#### (2) Derivation of the protein–ligand contact map

Here, the protein–ligand contact map is basically a matrix indicating the non-covalent interactions between each pocket residue and each ligand atom. During the supervised training process, PLANET was trained using real protein–ligand interactions observed in crystal structures as labels and learned to predict the protein–ligand contact map based on its input. The predicted protein–ligand contact map is helpful for rescaling the estimated partial energy contribution in the task of binding affinity prediction.

#### (3) Derivation of the intra-ligand distance matrix

The intra-ligand distance matrix includes the distance of all atom pairs within the ligand structure. Derivation of this matrix is an auxiliary task to help PLANET extract 3D features from the input 2D molecular graph.

In order to fulfill these three training objectives, PLANET comprises several modules, including protein feature extraction, ligand feature extraction, protein– ligand communication, ligand conformation prediction, pair-wise protein–ligand interaction prediction, and affinity prediction (Figure 1).

**Figure 1.**
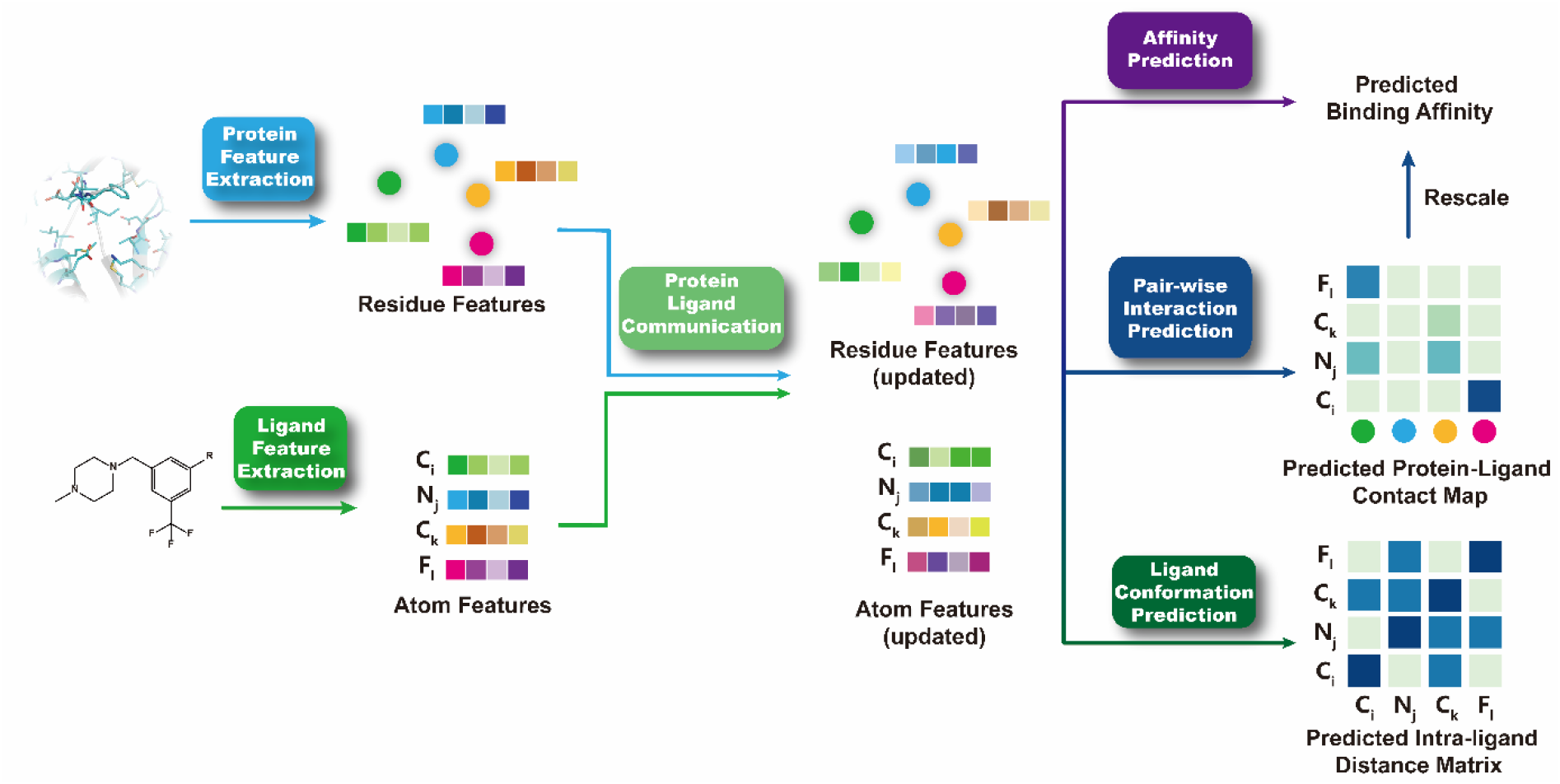
Illustration of the basic framework of PLANET.

#### Protein Feature Encoding and Extraction Module

The structure of a protein pocket is represented as an implicit graph in 3D space, in which each node denotes one pocket residue. The protein feature extraction module mainly consists of the Equivariant Graph Convolutional Layer (EGCL). Our implementation of EGCL is modified from the original Equivariant Graph Neural Network (EGNN) described by Satorras et al.^42^. EGNN was chosen by us because it has proven to be translationally, rotationally, and reflectionally equivariant and computationally more efficient than other models that require spherical harmonics. Given a protein–ligand complex, the ligand centroid of mass is calculated to define a sphere with a radius of 12.0 Å, within which all residues, except for non-natural residues, are considered as pocket residues and defined as nodes in the implicit graph. The initial feature for each node 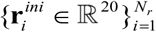 is the corresponding vector in the BLOSUM62 matrix ^43^ and the location for each node 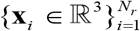 is the coordinate of the α-carbon in the corresponding pocket residue. *N_r_* denotes the number of pocket residues in the graph. Then, the initial features are transformed through an embedding layer (Equation 1):

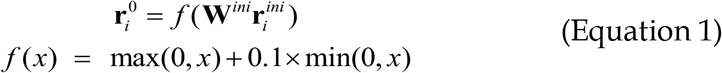

where *f* represents a LeakyReLU activation function and *f* shares the same definition hereafter. **W**^*ini*^ represents a single linear layer with learnable parameters that transforms the input features to a latent space of 300 dimensions. Hereafter, **W** shares the same definition, if not specified.

All nodes are updated through EGCL for several iterations. To reduce the number of parameters, the corresponding layers in all updating iterations share the same parameters (Equation 2):

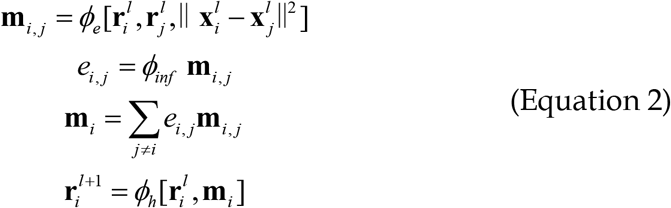

where *ϕ_e_* denotes two consecutive linear layers with two LeakyReLU activation functions and *ϕ_h_* denotes two consecutive linear layers with one LeakyReLU activation function. *ϕ_inf_* denotes one linear layer with one sigmoid activation function to normalize the output to (0, 1) as the relation between the two nodes. [,] represents the concatenation of vectors. After L_1_ update iterations (here, L_1_ = 3), the features of all pocket residues 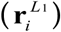 are obtained.

#### Ligand Feature Encoding and Extraction Module

The structure of a ligand is represented as a directed graph in 2D space, where each node denotes an atom and each edge denotes a covalent bond between the two corresponding atoms. The initial features are one-hot encoded descriptors of atoms 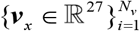 and bonds {***b***_*x,y*_ ∈ ℝ^6^} (See Table S1 in the *Supporting Information*). The ligand feature extraction module is modified from the graph encoder module in JT-VAE ^44^ by introducing a multi-head attention mechanism to better describe the contributions of different kinds of neighbors. Similar to the protein feature extraction module, layers share the same parameters in all update iterations. After L_2_ update iterations (here, L_2_ = 10), the features for all atoms 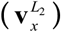 are obtained (Equation 3):

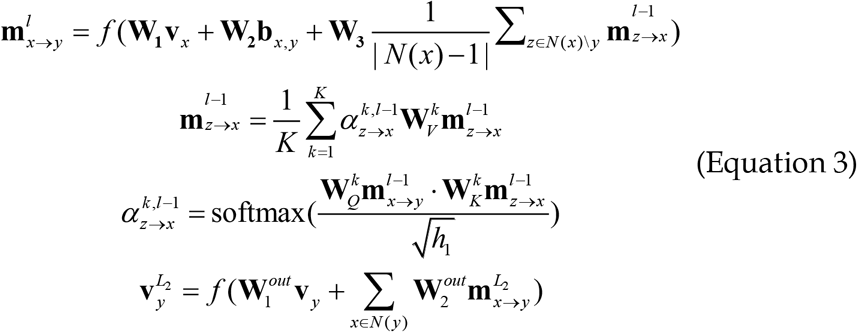

where | *N*(*x*) | denotes the number of neighbors of atom *x*, *N*(*x*) \ *y* denotes all neighbors of atom x except atom y. Dot (·) denotes the inner product of the matrix. K denotes the number of heads in the multi-head attention mechanism. *h*_1_ is the dimension of latent space (here, *h*_1_ =300). 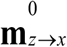 is zero-initialized.

#### Protein–Ligand Communication Module

In the aforementioned modules, protein and ligand features are updated in two independent graphs. To facilitate message exchange between protein and ligand features, a protein–ligand message exchange module was designed. The relation between each pair of pocket residue node and ligand atom node is derived in a cross-attention manner and served as weight in following feature update (Equation 4):

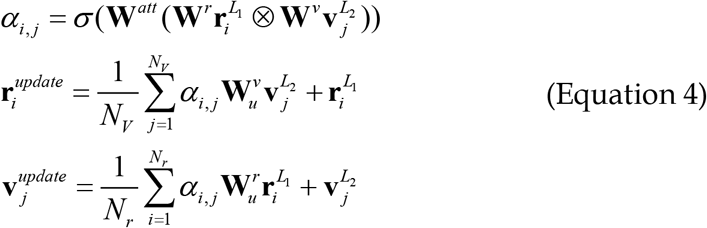

where N_v_ denotes the number of atoms in the molecule, σ denotes a sigmoid activation function, and ꕕ denotes element-wise production. For simplicity, the updated residue features 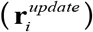 and atom features 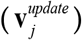 are abbreviated as **r**_*i*_ and **v**_*j*_, respectively.

#### Ligand Conformation Prediction Module

This module predicts how atoms interact with each other in 3D space. The obtained atom features (**v**_*x*_) are transformed to a suitable space to predict the interaction between the two atoms 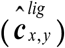. Distances (in angstroms) between each pair of atoms are calculated to set labels for the intra-ligand distance matrix according to the following criterion (Equation 5):

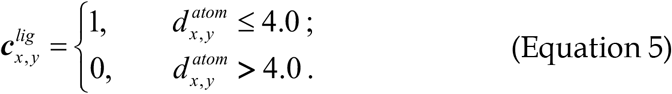

Binary cross entropy loss between the real label 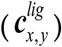 and the predicted value 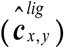 is computed as the loss of the ligand conformation prediction module (Equation 6):

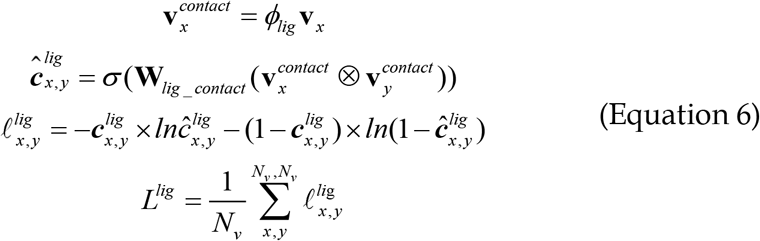

where **W**_*lig*_*contact*_ transforms the hidden features to one dimensional vector and *ϕ_lig_* is a single linear layer followed by a LeakyReLU activation function.

#### Protein–Ligand Interaction Prediction Module

The final residue features (**r**_*i*_ and atom features (**v**_*j*_) are transformed to predict the interaction 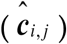 between each residue–atom pair. Labels for the protein–ligand contact map can be calculated from the crystal structure according to the following rule: if the distance of atom i to any atom in residue j is less than 4.0 Å, the label (***c***_*i*, *j*_) for atom i and residue j is set to 1; else, the label is set to 0 (Equation 7):

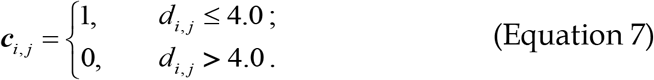

For a protein–decoy pair, all interaction labels are set to 0, indicating that such a protein and decoy molecule cannot form a stable complex. Binary cross entropy loss between the real label (***c***_*i,j*_) and predicted value 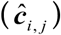 is computed as the loss (*L^pl^*) of the protein–ligand interaction prediction module (Equation 8):

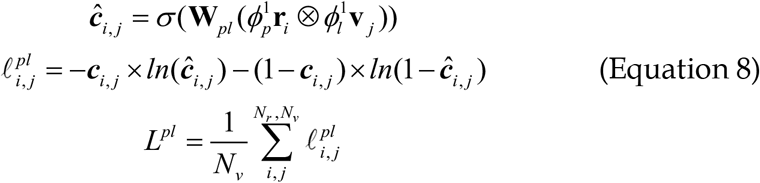

where **W**_*pl*_ transforms the hidden features to one dimensional vector. 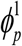 and 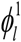 are two consecutive linear layers with two LeakyReLU activation functions.

#### Binding Affinity Prediction Module

According to the principles of physical chemistry, there is a linear relationship between protein-ligand binding free energy and negative logarithm of protein-ligand dissociation constant *K*_d_, i.e. −log K_d_ or p*K*_d_. We assume that the final residue features **r**_*i*_ and atom features **v**_*j*_ can be transformed to predict the partial energy contribution. It is reasonable to assume that in the whole protein-ligand complex, only the pocket residues and ligand atoms in direct interaction make contributions to binding energy. However, the interacting pairs of pocket residues and ligand atoms cannot be directly inferred from the input binding pocket and 2D ligand structure. To tackle this problem, the predicted protein-ligand interaction 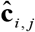 given by the protein-ligand interaction prediction module can be viewed as an interacting probability of each residue-atom pair, which is used to reweight the corresponding predicted partial energy contribution. All rescaled partial contributions are summed and the optimization objective is to minimize the L_2_ loss between the predicted affinity and experimental value (Equation 9):

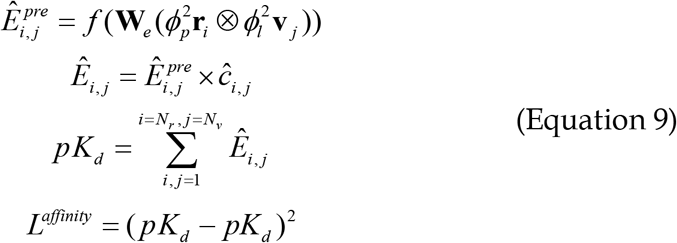

where **W**_*e*_ transforms the hidden features to one dimensional vector. 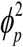 and 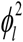 are two consecutive linear layers with two LeakyReLU activation functions.

#### Loss Function

The overall loss comes from the three above-mentioned prediction modules (Equation 10):

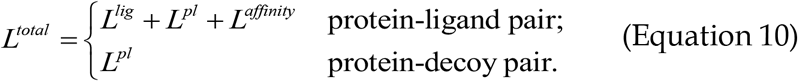

*L^lig^* and *L^affinity^* for a protein–decoy pair with unknown ligand conformation and binding affinity will be masked in the training process. It is not necessary to assign the binding affinity of a protein–decoy pair to a small value, since we have set the protein– ligand contact map labels to zero for a protein–decoy pair and therefore the partial energy contribution from each pair of pocket residue and ligand atom will be automatically rescaled to a small value, denoting for a low binding affinity. The scale of the three objectives were close to 10.0 at the beginning of training and rough adjustments on weights of these objectives only have minor impact on training. Thus, we gave equal weights on all objectives in the final version.

#### Model Implementation

PLANET was implemented using the PyTorch deep learning framework ^45^ along with the RDKit package ^46^ for chemical information processing.

### 2.2 Model Training

#### (1) Structural preparation

The PDBbind “general set” is essentially a large collection of protein-ligand complexes with available 3D structures from the Protein Data Bank (PDB) and corresponding experimental binding affinity data (i.e., K_d_, K_i_, and IC_50_) curated from literature 47. This data set has been a popular choice for deriving deep/machine learning models aiming at protein-ligand binding affinity prediction (Table 1). Our data set for training PLANET was also compiled by taking the PDBbind general set (v.2020), a total of 19443 protein-ligand complexes, as the starting pool. For each protein–ligand complex in this data set, structure preparation was performed using the *protein preparation* module in the Schrödinger software (v.2020), including fixing all incomplete residues, setting protonated states, and adding hydrogen atoms. The protonation state at pH 6.5~7.5 was set for the ligand molecule by using the *Epik* module ^48^. The resulting protein structure was saved in the PDB format, and the ligand structure was saved in the SDF format. A number of ligand structures could not pass the sanitize operation in RDKit and thus were removed. As result, a total of 19386 protein-ligand complexes in the PDBbind general set (v.2020) were successfully processed through the above procedure.

**Table 1.**
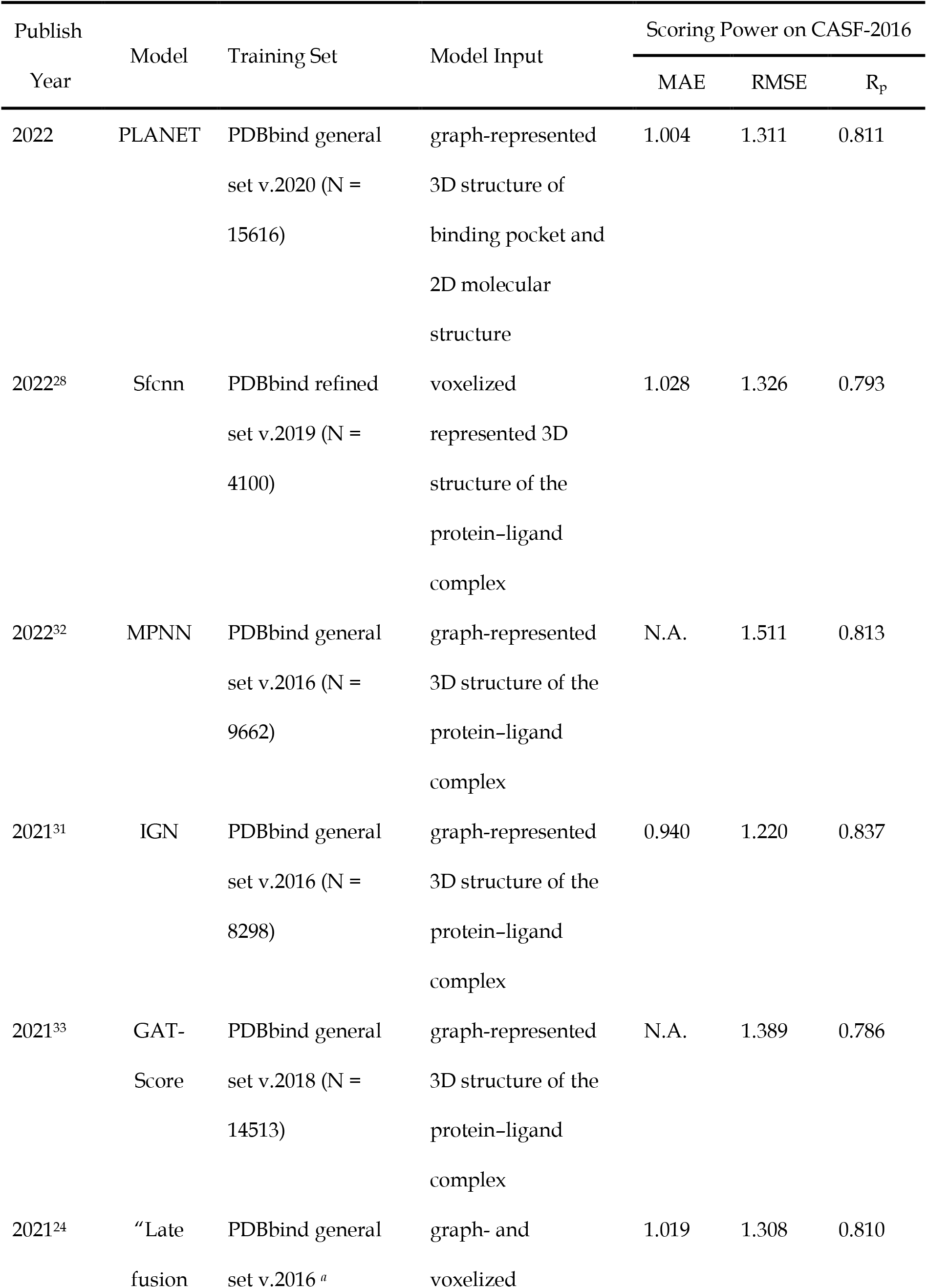
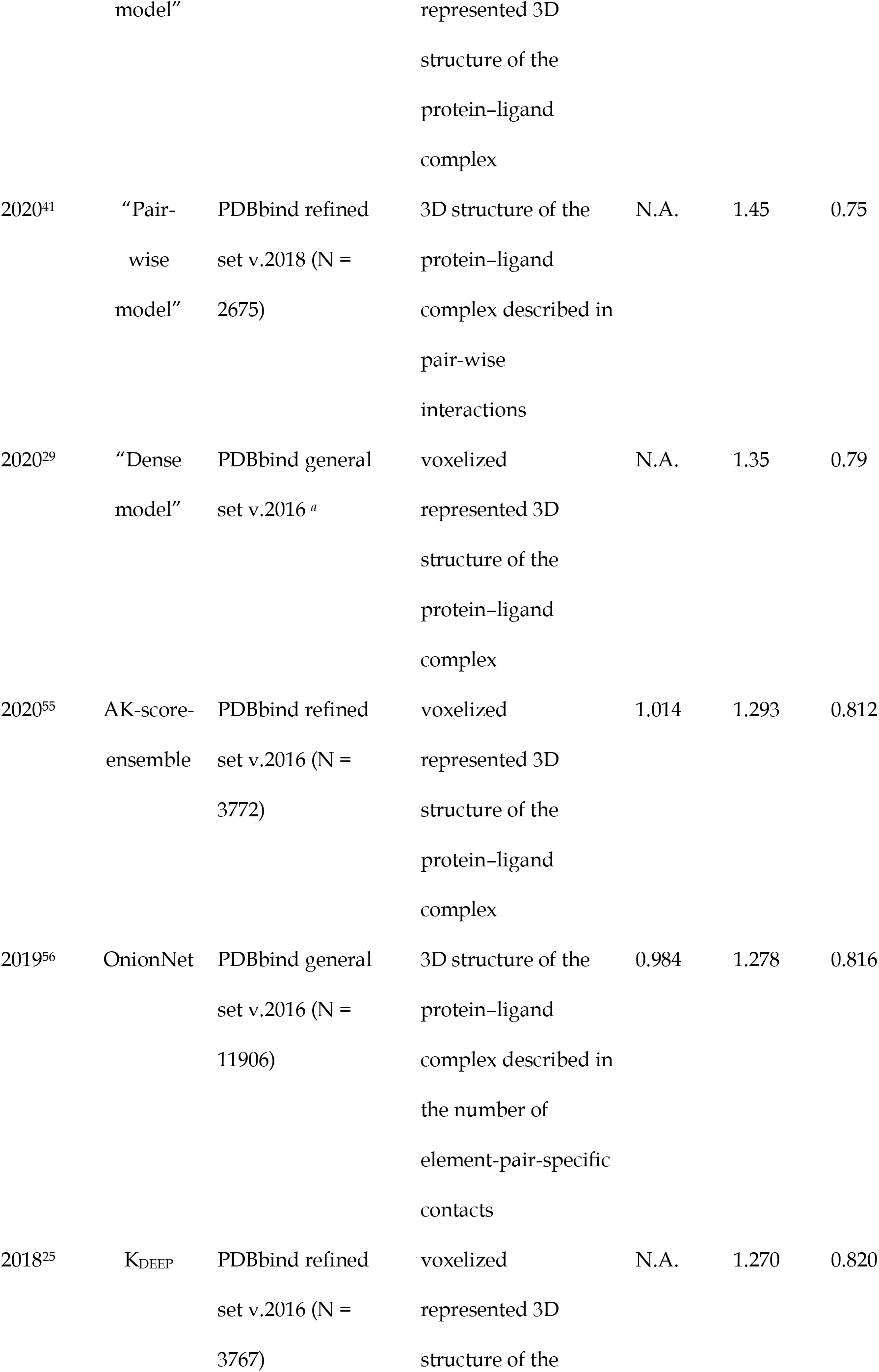
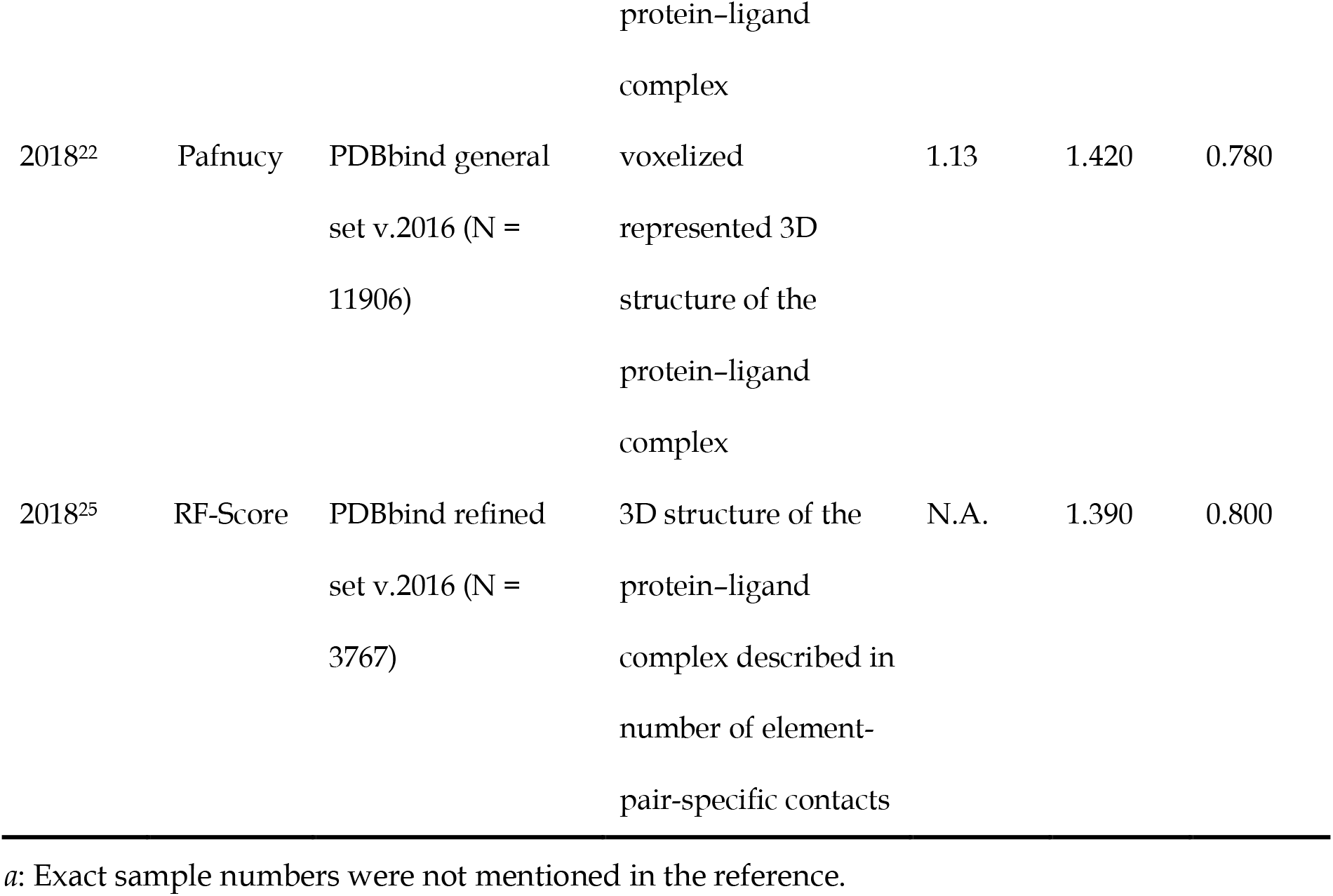
Basic information of some ML/DL models and their scoring power obtained on the CASF-2016 benchmark

| Publish<br>Year | Model | Training Set | Model Input | Scoring Power on CASF-2016 |  |  |
| --- | --- | --- | --- | --- | --- | --- |
|  |  |  |  | MAE | RMSE | R<sub>p</sub> |
| 2022 | PLANET | PDBbind general<br>set v.2020 (N =<br>15616) | graph-represented<br>3D structure of<br>binding pocket and<br>2D molecular<br>structure | 1.004 | 1.311 | 0.811 |
| 2022<sup>28</sup> | Sfcnn | PDBbind refined<br>set v.2019 (N =<br>4100) | voxelized<br>represented 3D<br>structure of the<br>protein-ligand<br>complex | 1.028 | 1.326 | 0.793 |
| 2022<sup>32</sup> | MPNN | PDBbind general<br>set v.2016 (N =<br>9662) | graph-represented<br>3D structure of the<br>protein-ligand<br>complex | N.A. | 1.511 | 0.813 |
| 2021<sup>31</sup> | IGN | PDBbind general<br>set v.2016 (N =<br>8298) | graph-represented<br>3D structure of the<br>protein-ligand<br>complex | 0.940 | 1.220 | 0.837 |
| 2021<sup>33</sup> | GAT-Score | PDBbind general<br>set v.2018 (N =<br>14513) | graph-represented<br>3D structure of the<br>protein-ligand<br>complex | N.A. | 1.389 | 0.786 |
| 2021<sup>24</sup> | “Late<br>fusion | PDBbind general<br>set v.2016 <sup>a</sup> | graph- and<br>voxelized | 1.019 | 1.308 | 0.810 |
|  | model" |  | represented 3D<br>structure of the<br>protein-ligand<br>complex |  |  |  |
| 2020<sup>41</sup> | "Pair-<br>wise<br>model" | PDBbind refined<br>set v.2018 (N =<br>2675) | 3D structure of the<br>protein-ligand<br>complex described in<br>pair-wise<br>interactions | N.A. | 1.45 | 0.75 |
| 2020<sup>29</sup> | "Dense<br>model" | PDBbind general<br>set v.2016 <sup>a</sup> | voxelized<br>represented 3D<br>structure of the<br>protein-ligand<br>complex | N.A. | 1.35 | 0.79 |
| 2020<sup>55</sup> | AK-score-<br>ensemble | PDBbind refined<br>set v.2016 (N =<br>3772) | voxelized<br>represented 3D<br>structure of the<br>protein-ligand<br>complex | 1.014 | 1.293 | 0.812 |
| 2019<sup>56</sup> | OnionNet | PDBbind general<br>set v.2016 (N =<br>11906) | 3D structure of the<br>protein-ligand<br>complex described in<br>the number of<br>element-pair-specific<br>contacts | 0.984 | 1.278 | 0.816 |
| 2018<sup>25</sup> | K<sub>DEEP</sub> | PDBbind refined<br>set v.2016 (N =<br>3767) | voxelized<br>represented 3D<br>structure of the protein-ligand<br>complex | N.A. | 1.270 | 0.820 |
| 2018<sup>22</sup> | Pafnucy | PDBbind general<br>set v.2016 (N =<br>11906) | voxelized<br>represented 3D<br>structure of the<br>protein-ligand<br>complex | 1.13 | 1.420 | 0.780 |
| 2018<sup>25</sup> | RF-Score | PDBbind refined<br>set v.2016 (N =<br>3767) | 3D structure of the<br>protein-ligand<br>complex described in<br>number of element-pair-specific contacts | N.A. | 1.390 | 0.800 |

*a*: Exact sample numbers were not mentioned in the reference.

#### (2) Compilation of the training set

Since our model was later evaluated on the CASF-2016 benchmark, the 285 protein-ligand complexes included in the CASF-2016 test set need to be removed from our data set to avoid overlap. Besides, we have demonstrated in a previous study that the existence of structurally similar samples between the training set and the test set, i.e. “soft overlap”, can lead to an over-optimistic performance of the resulting machine learning model ^49^. In order to remove the soft overlap from our training set, we employed the CD-hit software ^50, 51^ to cluster the entire PDBbind general set (v.2020) by protein sequence with a similarity cutoff of 90%. A protein-ligand complex was regarded as a redundant sample if (*i*) its protein sequence fell in the same cluster as any complex in the CASF-2016 test set, AND (*ii*) ECFP4 fingerprints of the ligand in this complex and any ligand in the CASF-2016 test set had a Tanimoto coefficient above 0.90. Consequently, a total of 369 complexes were identified as redundant and thus removed from our data set. After this step, the remaining complexes in each cluster were randomly split into two groups in a ratio of 80:20, where the first group was assigned to the training set and the second group to the validation set. In the end, our training set consisted of 15616 protein-ligand complexes, and the validation set consisted of 3401 protein-ligand complexes.

#### (3) Augment of non-binder decoys into the training set

The ligand molecule in a protein-ligand complex is of course regarded as a “binder”. In order to derive a model capable to differentiate binders from non-binders, it is a common practice to add some non-binders to the training set. In our work, a certain number of molecules with similar physicochemical properties as the ligand molecule in each protein-ligand complex in the training set were selected from ChEMBL release 29 (https://www.ebi.ac.uk/chembl/) ^52^ and then added to the training set as non-binder decoys. The workflow of our decoy selection is illustrated in Figure S1 in the *Supporting Information*. Briefly, molecular properties such as molecule weight, log*P*, number of hydrogen bond donor, number of hydrogen acceptor and so on were calculated using the RDKit software for all 2.1 million molecules in ChEMBL 29. For each protein-ligand complex in our training set, up to 100 candidate decoys were randomly selected from ChEMBL 29, and those met at least six of the following eight criteria were selected out: (1) Difference in molecule weight was less than 10%; (2) Difference in log*P* is less than 1 units; (3) Difference in number of hydrogen bond donor was less than 1; (4) Difference in number of hydrogen bond acceptor was less than 1; (5) Difference in number of chiral centers was less than 1; (6) Difference in number of rings was less than 1; (7) Difference in number of heavy atoms was less than 1; (8) Difference in number of total atoms was less than 1. Note that the qualified candidates were further examined by docking them into the same binding site as the ligand molecule in the given protein-ligand complex. Here, molecular docking was conducted by using the Glide SP module in the Schrödinger software (v.2020) with default settings, except for “INNERBOX” set to 5 Å and “GLIDE_CONFGEN_EFCUT” set to 4.0 kcal/mol. The candidates for which Glide SP failed to generate any valid binding pose were finally accepted as the non-binder decoys for the given protein-ligand complex. This additional molecular docking examination was conducted to reduce the chance that the selected non-binder decoys were actually binders. As result, a total of 161547 non-binder decoys were added to our training set to augment the 15616 binders. Each binder was thus accompanied by roughly 10 non-binder decoys.

#### (4) The training process

A batch size of 16 samples was used during training, where the chance of retrieving a true protein-ligand complex or a protein-decoy pair as one sample was equal. The training process was performed with an Adam optimizer with an initial learning rate of 0.0001 for updating parameters. Model parameters were saved every 5000 batches. The whole training process was completed in 1.5 days on a single NVIDIA P100 GPU card with 16 GB memory. The model with the best performance on the validation set was selected for further evaluation.

### 2.3 Model Evaluation

#### (1) Evaluation of scoring power

Following the terminology in this field, scoring power refers to the ability of a scoring model to predict protein-ligand binding affinity based on its input. In our work, the scoring power of PLANET was tested on the CASF-2016 benchmark ^10, 47^. CASF-2016 has been a well-established benchmark for testing scoring functions, including various machine learning models (see Table 1 for some examples). The primary test set in CASF-2016 consists of 285 diverse protein–ligand complexes with high-quality structures and experimental binding affinity data that are selected from the PDBbind general set (v.2016). In this test, evaluation metrics computed by us including the mean absolute error (MAE), root mean square error (RMSE), Pearson correlation coefficient (R_p_), and Spearman correlation coefficient (R_s_).

PLANET was also evaluated regarding its performance in predicting intra-ligand distance matrix and protein–ligand contact map. For each task, its performance was measured by the cosine similarity between the real values and predicted ones on the entire CASF-2016 test set. In addition, area under the curve of receiver operating characteristic (AUC ROC) and area under the curve of precision-recall (AUC PR) were calculated at the atom level, residue level, and pairwise level in the task of protein– ligand contact map prediction ^37^. To calculate AUC ROC and AUC PR values at the atom level, ligand atoms with at least one contact with any pocket residue were labeled as positive samples. The predicted value for a ligand atom was the sum of all contacts formed between it and the protein. Cases in which all the ligand atoms were positive samples were excluded from AUC ROC and AUC PR calculations. The same method was also applied to the calculation of AUC ROC and AUC PR values at the residue level.

In order to testify the predictive power of PLANET further, a random baseline was added to the comparison. This model was designed to generate intra-ligand distance matrix and protein–ligand contact map randomly by referring to the true ligand structure and protein-ligand complex structure, respectively. For example, assuming that a given protein-ligand complex structure is projected into a protein-ligand contact map consisting of 95% false values (0) and 5% true values (1), this baseline model would fill up a predicted protein-ligand contact map with 95% values in the range of (0.5, 1.0) and 5% values in the range of (0, 0.5) in a totally random manner. This baseline random model was also applied to the entire CASF-2016 test set. Its performance was measured by the same set of evaluation metrics applied to PLANET.

#### (2) Evaluation of screening power

In this article, the “screening power” refers to the ability of a scoring model to differentiate binders from non-binders. Two public virtual screening benchmarks, DUD-E ^39^ and LIT-PCBA ^40^, were employed in our work to test the screening power of PLANET. DUD-E contains 102 target proteins, where each target protein is associated with (i) a protein structure along with a known ligand to define the binding pocket, and (ii) relevant binders as well as non-binders in a ratio of roughly 1:50. LIT-PCBA contains 15 target proteins, where each target protein is associated with (i) a certain number of protein–ligand complex structures, and (ii) relevant binders as well as non-binders in a ratio of roughly 1:1000. All protein structures and binder and non-binder molecules in these two benchmarks were prepared following the same procedure for processing the PDBbind general set. Note that because LIT-PCBA provides multiple protein-ligand complexes for each target protein, we only considered the one with the highest resolution in order to save computational cost (see Table S2 in the *Supporting Information*).

In this test, the AUC ROC score and enrichment factor (EF) at different thresholds (Equation 11) were computed as the measurements of screening power. Enrichment factor was computed with the following formula:

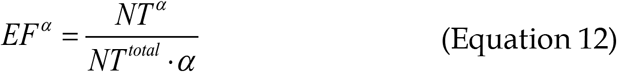

where *NT*^α^ is the number of binders found among the top α-ranked molecules (e.g., α = 1%) based on the predicted binding affinity. *NT*^total^ is the total number of binders for the given target in the data set.

To make a comparison to conventional docking method, the Glide SP module in the Schrödinger software was also tested on the LIT-PCBA benchmark. Glide SP was applied all with default settings. Note that Glide SP failed to generate any docking pose for some molecules on certain target proteins. The docking scores for the molecules in such cases were assigned to zero for the sake of convenience.

### 2.4 Projection and Visualization of Data Sets

The uniform manifold approximation and projection (UMAP) analysis ^53^ was used to illustrate the chemical space covered by the several data sets employed in our work. For this purpose, all ligand molecules (i.e. the binders) and the selected decoy molecules (i.e. non-binders) in the PDBbind general set and all binders and 20% of the non-binders in the DUD-E and LIT-PCBA data set were gathered. The ECFP4 fingerprints of all gathered molecules were calculated and hashed to 512 bits using the RDKit software, resulting in binary vectors with 512 dimensions. Those vectors were then subjected to the incremental principal component analysis reduction ^54^ to reduce their dimension to 100. Finally, UMAP analysis was conducted using the *umap-learn* python package with default settings to project the reduced vectors to two dimensions for visualization.

## 3. Results and Discussion

### 3.1 Prediction of Protein–Ligand Binding Affinity

As explained in the Model Architecture section, PLANET was trained in a multi-objective manner, i.e. predicting protein–ligand contact map, intra-ligand distance matrix, and protein–ligand binding affinity. The primary goal of PLANET from a practical perspective is to predict protein–ligand binding affinity. Following the convention in this field, we first tested its “scoring power”, i.e. computing the binding affinity of a protein-ligand complex based on its structure, on the CASF-2016 test set, which consists of 285 diverse protein–ligand complexes. The correlation between the experimental binding affinity data and the predicted valued by PLANET is shown in Figure 2, where the Pearson correlation coefficient is 0.811 and the RMSE is 1.311 log units. A comparison of the basic information, including the scoring power obtained on the same benchmark, of a number of deep/machine learning models published in recent years is summarized in Table 1. Most of these models are able to produce a Pearson correlation coefficient around 0.80 on the same test set. PLANET is at least close to the best of these in terms of its statistical metrics.

**Figure 2.**
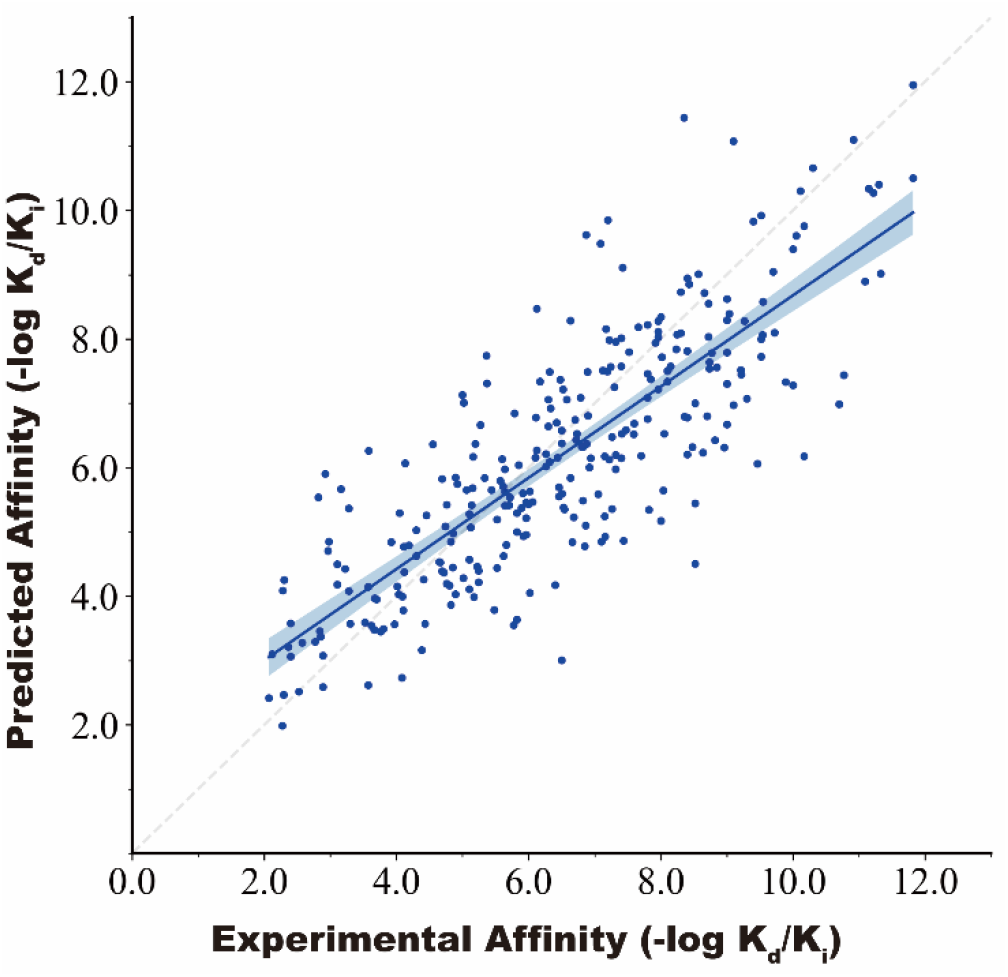
Correlation between the experimental binding affinity data and the predicted values by PLANET on the CASF-2016 test set (N = 285; R_p_ = 0.811; RMSE = 1.311 log units). The solid line is the regression line, where the 95% confidence interval is indicated by the shaded region.

Here, the readers should be reminded that the real “scoring power” of PLANET may be underrated by these statistical metrics for two reasons. First, all other models included in Table 1 directly take 3D protein–ligand complex structures as input, while PLANET only uses 3D binding pocket structures and 2D ligand structures. Obviously, it is more challenging to predict protein–ligand binding affinity without the exact information of their binding mode. In fact, the scoring power of some deep learning models that rely on crystal protein–ligand complex structures would decrease considerably if complex structures derived from molecular docking are used instead. In the case of the “fusion model” reported by Jones et al. ^24^, for example, when complex structures derived from molecular docking were used instead of crystal structures, the R_p_ and RMSE values of their model dropped from 0.810 and 1.308 log units to 0.712 and 1.871 log units, respectively, on the CASF-2016 test set. We suspect that other deep/machine learning models included in Table 1 would exhibit a lower scoring power if they were subjected to the same treatment. The scoring power test enabled by the CASF-2016 benchmark is certainly a basic test of the essential quality of a protein–ligand interaction scoring function. However, in reality, it is not very meaningful to “predict” the binding affinity of a protein–ligand pair based on their crystal complex structure because it is far more difficult to obtain crystal complex structures than measuring binding affinity experimentally. From this perspective, our PLANET model is more robust and more efficient for the purpose of protein–ligand binding affinity prediction without the need of obtaining or generating the corresponding protein-ligand complex structures in prior.

Second, during model training, we removed the protein–ligand complexes that are structurally too similar to those in the CASF-2016 test set with a similarity cutoff of 90% applied to the protein and ligand sides. We call such samples “soft overlap” between the training set and the test set. We have demonstrated in our previous study ^49^ that the existence of such samples in the training set leads to over-optimistic statistical metrics in the scoring power test for machine learning scoring functions. Therefore, we suggested that removing the soft overlap between the training and test sets should be a standard practice for testing a deep/machine learning model. However, none of the models listed in Table 1 actually executed this standard practice in their training process. It is unclear to what extent their statistical metrics (R_p_ and RMSE) in this scoring power test would decrease if they did so.

A special feature in the working mechanism of PLANET needs to be emphasized here. As explained in the Method section, PLANET predicts binding affinity by integrating only the contributions from the interacting pairs of pocket residues and ligand atoms. This mechanism requires PLANET to derive the protein-ligand contact map at the first place. This, however, is not a straightforward task because the inputs to PLANET from the ligand side are merely 2D molecular graphs. Thus, prediction of the protein-ligand contact map was included as a major objective in our model training. Besides, an auxiliary objective to derive the intra-ligand distance matrix was added to help PLANET extract 3D features from 2D molecular graphs. Such a working mechanism is quite different from many deep learning models listed in Table 1, which typically rely on 3D protein-ligand complex structures for prediction. Importantly, inclusion of protein-ligand contact map and intra-ligand distance matrix predictions also makes our model interpretable, at least to some extents, in terms of capturing the essential 3D features in protein–ligand interaction.

The quality of the protein–ligand contact map and intra-ligand distance matrix predicted by PLANET is measured by the cosine similarity between the corresponding matrix derived from the real structure and the predicted matrix. The statistical results obtained on the entire CASF-2016 test set are summarized in Table 2. In addition, as prediction of protein–ligand contact map and intra-ligand distance matrix is essentially a binary classification task, AUC ROC and AUC PR values at different levels were also calculated and summarized in Table 2. Generally, PLANET is able to predict the intra-ligand distance matrix well with an overall cosine similarity of 0.926 on the CASF-2016 test set. In comparison, the overall cosine similarity given by the baseline model is 0.518 in the same case. Regarding protein–ligand contact map prediction, the overall cosine similarity given by PLANET is 0.587. This result seems to be somewhat less promising. But considering the overall cosine similarity given by the baseline model is as low as 0.200 in this case, one can conclude that the accuracy of PLANET in this aspect is still encouraging. Besides, as measured by the AUC ROC and AUC PR values calculated at different levels, the accuracy of our PLANET is superior to that of the baseline model by a large margin in both intra-ligand distance matrix and protein-ligand contact map prediction (Table 2). As discussed earlier in this article, we believe that the predictive power of PLANET is rooted in its capability of capturing the essential 3D features in protein-ligand interactions. To provide some explicit examples, the predicted intra-ligand distance matrices and protein–ligand contact maps for two selected protein–ligand complexes in the CASF-2016 test set are illustrated in Figure 3. One can see that the predicted matrices largely resemble the ones derived from real structures.

**Table 2.** Performance of PLANET and a baseline model in intra-ligand distance matrix and protein–ligand contact map prediction on the CASF-2016 test set

| Tasks |  | Cosine Similarity | AUC ROC | AUC PR |
| --- | --- | --- | --- | --- |
| Performance of our PLANET model |  |  |  |  |
| Intra-ligand distance matrix | | 0.926±0.050 | 0.976±0.033 | 0.961±0.064 |
| Protein-ligand contact map | Atom-level | | 0.725±0.235 | 0.965±0.047 |
| | Residue-level | 0.587±0.120 | 0.956±0.052 | 0.934±0.069 |
| | Pairwise-level | | 0.932±0.044 | 0.517±0.173 |
| Performance of a baseline model |  |  |  |  |
| Intra-ligand distance matrix | | 0.518±0.104 | 0.500±0.017 | 0.384±0.128 |
| Protein-ligand contact map | Atom-level | | 0.513±0.193 | 0.908±0.090 |
| | Residue-level | 0.200±0.022 | 0.504±0.086 | 0.351±0.114 |
| | Pairwise-level | | 0.502±0.027 | 0.060±0.013 |

**Figure 3.**
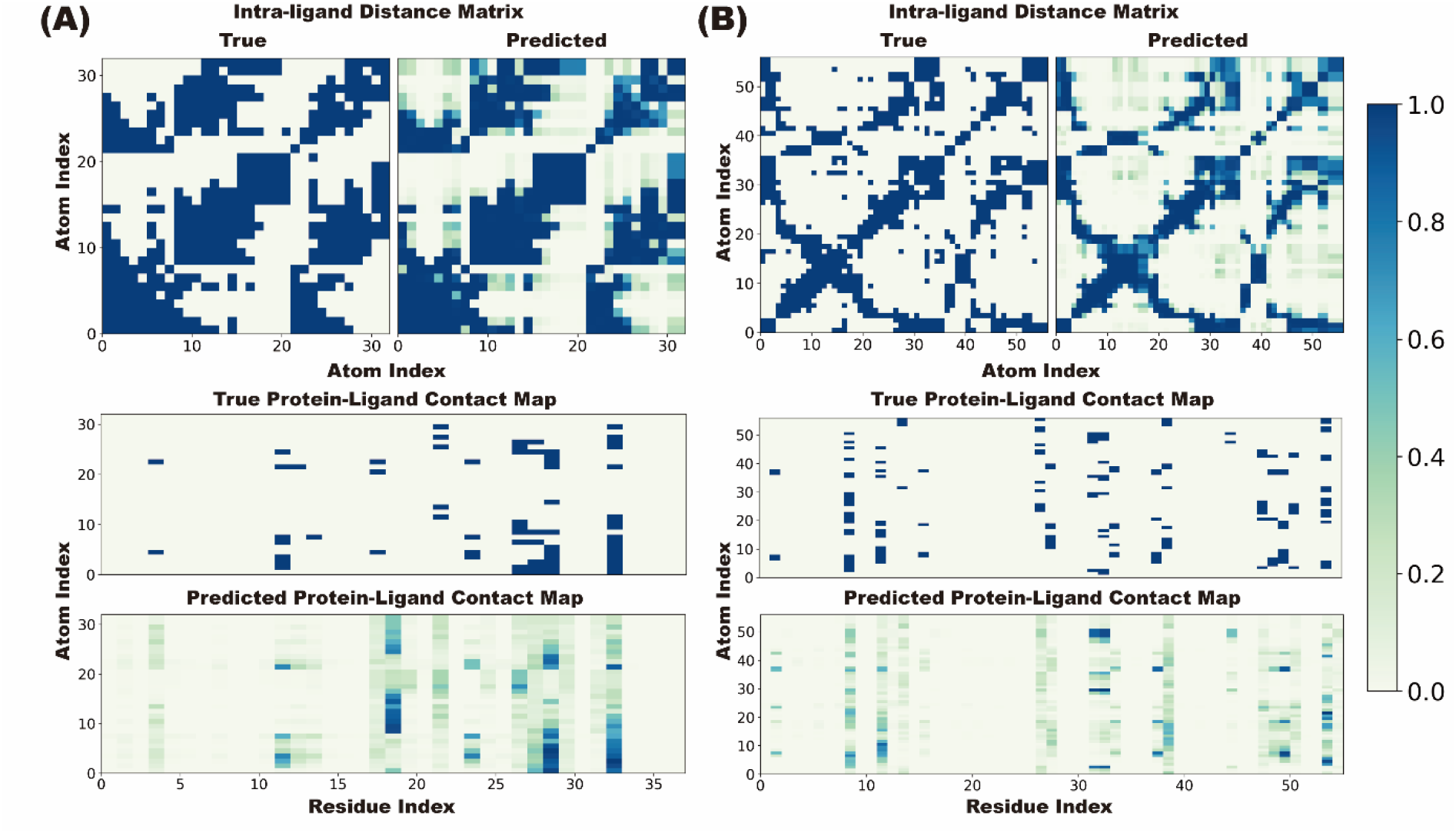
Visualization of the intra-ligand distance matrix and protein–ligand contact map predicted by PLANET. (A) PDB entry 3L7B: experimental −log Ki = 2.40, predicted value = 3.07; (B) PDB entry 5DWR: experimental −log Ki = 11.22, predicted value = 10.27.

As suggested by our evaluation results, accurate prediction of the protein–ligand contact map is a more challenging task for PLANET under its current framework. Protein–ligand contact map prediction is assigned as a binary classification task, but the labels in our training set are extremely unbalanced. In fact, the labels for more than 95% of possible residue–atom pairs are blank. Such unbalanced training data of course presented a troublesome challenge in our model development. Note that the AUC ROC and AUC PR values computed at the residue-level (0.956 and 0.934, respectively) indicate that PLANET captures most of the important residues interacting with the ligand molecule, albeit some false positives exist among the predictions. Nevertheless, our opinion is that more accurate prediction of the protein–ligand binding mode should be accomplished by other deep learning models in a different framework, such as the one described by Mendez-Lucio ^57^ and Shen ^58^.

### 3.2 Performance in Virtual Screening

It should be mentioned again that although the “scoring power” is an essential aspect of a successful scoring model, in reality scoring models are applied more often to virtual screening tasks, i.e. differentiating true binders from non-binders. In other words, the “screening power” of a scoring model should receive more attention. Several CNN models described in literature, such as Pafnucy and OnionNet, were shown to achieve higher scoring power on the CASF-2016 test set than traditional scoring functions, but they could hardly distinguish between binders and non-binders in virtual screening trials ^34^. A common strategy for achieving a good screening power through deep learning is to derive a classification model by using a special training set consisting of binders and non-binders ^31, 59–62^. This strategy is of course sound in concept. However, if the training set and the chosen test set are congeneric in nature, e.g. splitting a common mother data set ^49, 63^, deep learning models tend to exhibit over-optimistic performance.

In order to avoid this potential caveat in model evaluation, we tested the screening power of PLANET on the DUD-E benchmark directly after its training on the PDBbind data set was accomplished, i.e. no further fine-tuning was attempted to extend its generalizability. The results produced by PLANET, together with those produced by OnionNet, Pafnucy, and several other machine learning models reported by Shen et al. ^34^, are summarized in Table 3. One can see that the performance of all deep /machine learning models originally trained on the PDBbind data set were much less promising in this test, because (i) the composition of the DUD-E data set is distinctively different from the PDBbind data set, and (ii) the goal switches from regression to classification in this scenario. In contrast, our PLANET exhibited a reasonable level of performance here. For example, PLANET (AUC ROC = 0.761) is superior to Pafnucy (AUC ROC = 0.631) and OnionNet (AUC ROC = 0.597), given that all these three deep learning models were trained on the PDBbind data set and achieved a comparable level of scoring power on the CASF-2016 benchmark.

**Table 3.** Performance of some deep learning and machine learning models and conventional scoring functions on the DUD-E benchmark.

| Model | AUC ROC |  | EF0.5% | EF1% | EF5% |
| --- | --- | --- | --- | --- | --- |
| | mean±std | median | mean | mean | mean |
| PLANET | 0.761±0.123 | 0.770 | 10.234 | 8.832 | 5.402 |
| $\Delta_{\text{VinaRF}_{20}}$ <sup>a</sup> | 0.697±0.102 | 0.695 | 9.516 | 7.995 | 4.382 |
| RFscore-V4 <sup>a</sup> | 0.652±0.108 | 0.655 | 4.902 | 4.521 | 2.981 |
| NNscore2.0 <sup>a</sup> | 0.683±0.098 | 0.683 | 4.162 | 4.022 | 3.123 |
| Pafnucy <sup>a</sup> | 0.631±0.124 | 0.639 | 4.241 | 3.861 | 2.767 |
| OnionNet <sup>a</sup> | 0.597±0.102 | 0.606 | 2.840 | 2.840 | 2.205 |
| Glide SP <sup>a</sup> | 0.767±0.116 | 0.783 | 19.389 | 16.182 | 7.231 |
| Vina <sup>b</sup> | 0.716 | N.A. | 9.139 | 7.321 | 4.444 |

a: Performance metrics are cited from ref <sup>34</sup>.
b: Performance metrics are cited from ref <sup>60</sup>.

Very interestingly, the classical scoring function Glide SP achieved the best performance among all models in terms of AUC ROC score and EFs. This reflects the fact that although classical scoring functions are limited by their simple mathematical architecture in encoding a large, complex body of data, they are able to capture some basic physics in protein-ligand interaction and thus may have a generalized performance in virtual screening tasks. Another piece of evidence supporting this observation comes from the results of Δ_Vina_RF_20_, which also outperformed other deep/machine learning models in this test. Note that Δ_Vina_RF_20_ is in fact a combination of the Vina scoring function and a correction term derived through a Random Forest model 9. That correction term has improved the scoring power of Δ_Vina_RF_20_. However, as one can see in Table 3, it brings no advantage to the screening power as compared to Vina. In other words, the screening power of Δ_Vina_RF_20_ is actually established by the Vina scoring function. Although our PLANET model is second to Glide SP on the DUD-E benchmark, it outperformed all other deep/machine learning models. PLANET demonstrated a good level of generalizability even though the goal here has shifted from the original regression task.

Although DUD-E is perhaps the most popular benchmark for evaluating virtual screening models, a few studies have pointed out that the artificially generated decoys included in the DUD-E data set may lead to biased results ^40, 63–66^. In order to conduct a more objective evaluation of screening power, we further tested PLANET head-to-head with Glide SP on the recently published LIT-PCBA benchmark ^40^. In this new benchmark, all binders and decoys are experimentally proven according to the information from PubChem BioAssay ^67^. All 2.7 million molecules are carefully selected through an asymmetric validation embedding procedure. Besides, for each target included in LIT-PCBA, the number of decoys is roughly 1000 times larger than that of binders, which better mimics the real situation in virtual screening.

Similar to the case of DUD-E, our PLANET model was directly evaluated on LIT-PCBA without fine-tuning. Glide SP was applied in its standard workflow and parameter setting. The evaluation results are illustrated in Figure 4. More details can be found in Table S3 in the *Supporting Information*. Here, one can see that LIT-PCBA is indeed more challenging than DUD-E, as the mean AUC ROC scores of PLANET and Glide SP are 0.573 and 0.553, respectively, both of which are lower than the counterparts obtained on DUD-E. Compared to Glide SP, PLANET produced higher AUC ROC scores on 10 targets, comparable scores on two targets, and lower scores on three targets in LIT-PCBA. Nevertheless, paired *t*-test revealed no significant difference between the results produced by PLANET and Glide SP, i.e. *p* = 0.466 for AUC ROC scores and *p* = 0.248 for EF^1.0%^. Thus, the screening power demonstrated by these two models on the LIT-PCBA benchmark is basically on par. Nevertheless, PLANET had a huge technical advantage in terms of efficiency: It took PLANET about 3 hours to finish all jobs needed by this test on a single NVIDIA P100 GPU card; while it took Glide SP about 25 days to finish on an Intel XEON W-2245 CPU (16-processor at 3.90 GHz). PLANET achieves its computational efficiency by skipping the time-consuming molecular docking process and replaces it by prediction of the protein– ligand contact map. Thus, PLANET can hopefully become a practical tool for virtual screening of extremely large compound databases in addition to the current approaches ^68–71^.

**Figure 4.**
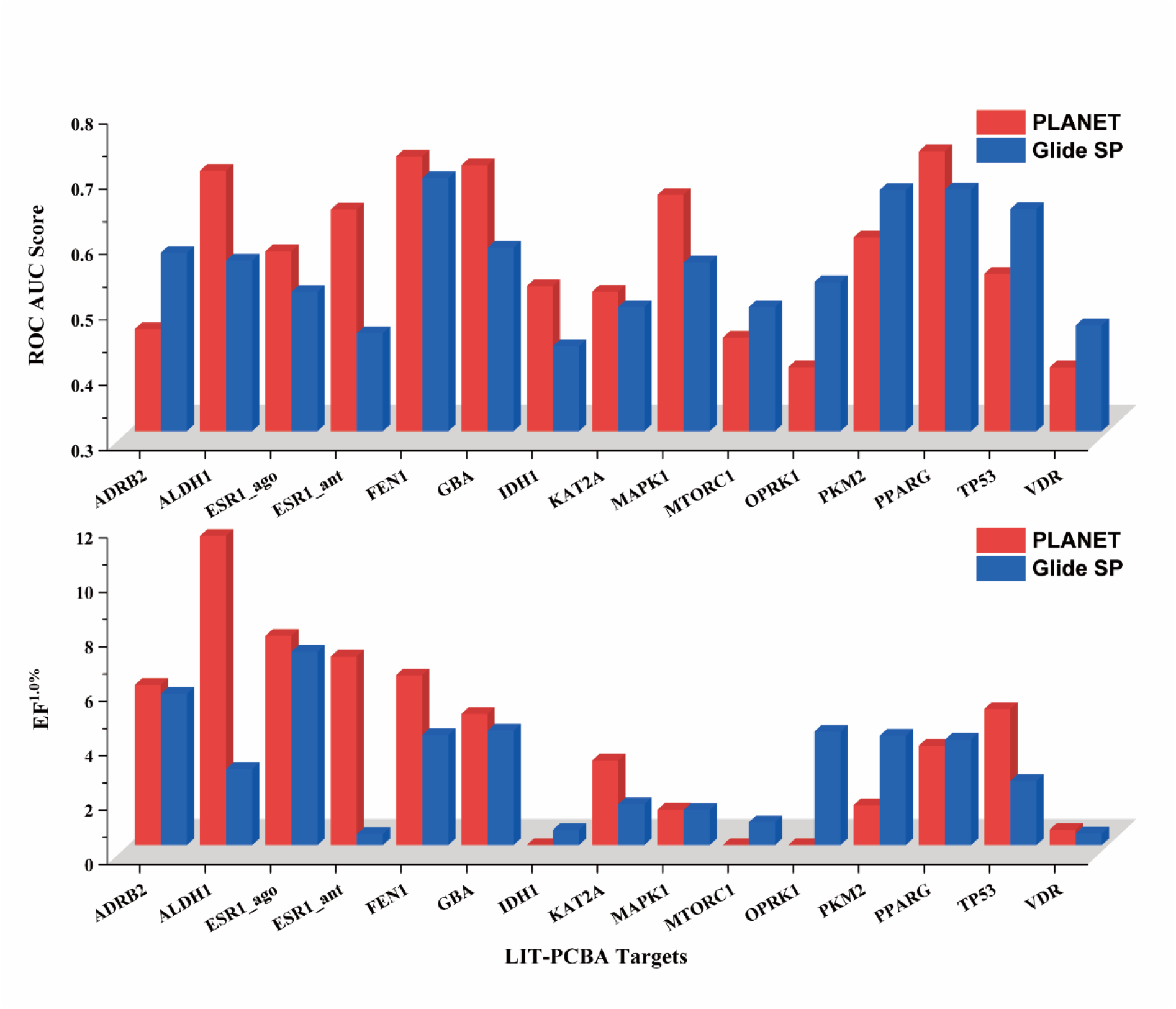
Evaluation results of PLANET and Glide SP on the LIT-PCBA data set.

We have noticed that Pafnucy, a CNN-based deep learning model, was also tested on the LIT-PCBA benchmark ^72^. According to the reported results, the screening power of Pafnucy was rather poor on this benchmark. In fact, nearly 90% of the binding affinities predicted by Pafnucy are distributed in a narrow range between 5.0 and 6.5 (in −log K_d_/K_i_ units) in a basically target-independent manner (Figure 5B). This failure may result from the voxelized representation of the protein binding site by a CNN model, which produces a sparse tensor with extremely high dimensions and it is in turn difficult to extract useful features from the sparse tensor ^63^. In contrast, PLANET produced binding affinity in a much wider range in a more realistic, target-dependent manner (Figure 5A). This suggests that PLANET does have some “understanding” of protein–ligand interactions and thus make meaningful predictions.

**Figure 5.**
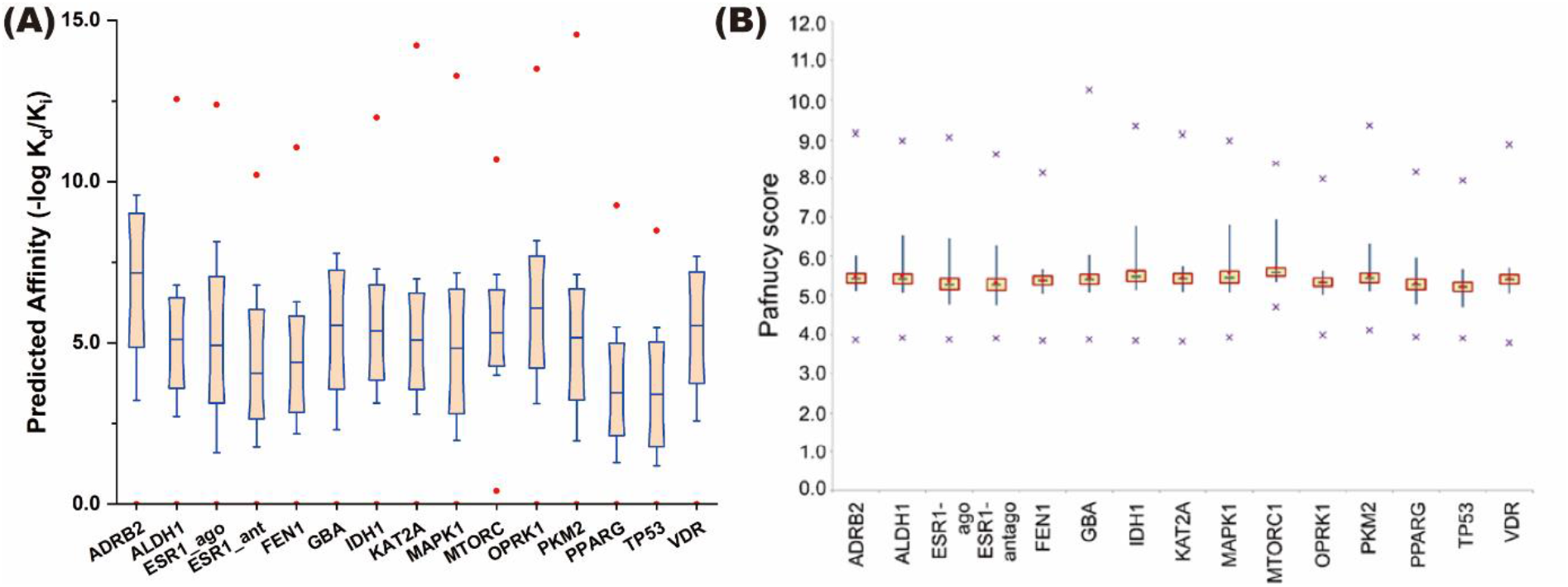
(A) Distribution of the predicted protein-ligand binding affinities given by PLANET on the LIT-PCBA data set. Labels on the x axis are the names of each target protein. The boxes delimit the 10th and 90th percentiles of the predicted values; while the whiskers delimit the 5th and 95th percentiles. The mean value is indicated by the blue dash in the middle of each box. The minimum and maximum values are indicated by red dots. (B) Distribution of the predicted protein-ligand affinities given by Pafnucy on the same LIT-PCBA data set. This figure is cited from Ref ^72^, which is rendered in a similar style as described above.

### 3.3 Impact of the Training Set on Model Performance

Besides predicting protein-ligand binding affinity, our PLANET model has additional training objectives including predicting intra-ligand distance matrix and protein–ligand contact map. Additional training data are thus needed for those objectives. In order to reveal the improvements achieved by using additional training objectives and training sets, we performed several ablation studies as follows: (1) “*PLANET_crystal*”: This model shared the same three objectives as the full model, but it was trained only on the crystal protein–ligand complex structures in the PDBbind data set, with no decoy molecules were added; (2) “*PLANET_2D*”: This model did not include the objective of predicting intra-ligand distance matrix, but its training set was the same as the full model; (3) “*PLANET_baseline*”: This model did not include the objective of predicting intra-ligand distance matrix, and it was trained only on the crystal protein–ligand complex structures in the PDBbind data set. The above three additional models were trained in a similar process as the full model. After training, they were tested on the CASF-2016 benchmark to evaluation their scoring power and on the DUD-E benchmark to evaluate their screening power. The evaluation results are summarized in Table 4.

**Table 4.** Scoring power and screening power of the trained models in several ablation studies.

| Model | Scoring Power |  |  |  | Screening Power |  |  |  |
| --- | --- | --- | --- | --- | --- | --- | --- | --- |
|  | As on CASF-2016 |  |  |  | As on DUD-E |  |  |  |
|  | MAE | RMSE | R<sub>p</sub> | R<sub>s</sub> | AUC | EF<sup>0.5%</sup> | EF<sup>1%</sup> | EF<sup>5%</sup> |
| PLANET | 1.005 | 1.311 | 0.811 | 0.811 | 0.761 | 10.234 | 8.832 | 5.402 |
| *PLANET_crystal* | 0.977 | 1.278 | 0.814 | 0.803 | 0.661 | 5.253 | 4.677 | 3.112 |
| *PLANET_2D* | 1.044 | 1.336 | 0.797 | 0.797 | 0.692 | 5.514 | 4.860 | 3.534 |
| *PLANET_baseline* | 1.000 | 1.260 | 0.820 | 0.806 | 0.651 | 4.828 | 4.284 | 2.904 |

As shown in Table 4, the scoring power of all three additional models was roughly at the same level, i.e. R*p* around 0.80 on the CASF-2016 test set. Interestingly, when the decoys were removed from our training set, the resulting model, i.e. *PLANET_crystal*, even achieved a marginally better soring power (MAE = 0.977 log unit) than PLANET itself (MAE =1.005 log unit). This indicates that the PDBbind data set itself is sufficient for training a model with a promising scoring power on CASF-2016, which is in fact a common practice for most of the deep/machine learning models summarized in Table 1. This is however not a very challenging task because the test set in CASF-2016 is also derived from the PDBbind data set, and thus the test set and the training set are homogeneous in this scenario.

As it comes to the screening power, substantial difference is witnessed among the several models. For example, the AUC ROC score produced by PLANET is 0.761 while the value by *PLANET_crystal* is 0.661. This gap between the performance of two models indicate that adding decoy molecules into the training set helps to build a better screening power. Note that the screening power refers to the ability to distinguish between binders and non-binders. The PDBbind data set is formed by known protein-ligand complexes, i.e. all ligand molecules included in this data set are strong or weak binders. Consequently, such a data set is not sufficient for deriving a deep learning model with a good screening power because the model has no chance to learn about any non-binder during the training process. In fact, the strategy of adding decoys into the training set has been applied by other deep learning models published in literature.^38^ However, our PLANET model has implemented one additional feature to further enhance its screening power, i.e. prediction of intra-ligand distance matrix. The improved performance of PLANET (AUC ROC = 0.761) over *PLANET_2D* (AUC ROC = 0.692) indicates that the task of predicting intra-ligand distance matrix encourages PLANET to extract features from 2D molecular graphs and in turn helps with its screening power. If disabling the task of intra-ligand distance matrix prediction and also removing decoys from the training set, it is not surprising to observe that the resulting model, i.e. *PLANET_baseline*, exhibited the lowest screening power (AUC ROC = 0.651).

Frankly speaking, the screening power of our PLANET model is still not satisfactory. For example, its performance on either DUD-E or LIT-PCBA has not surpassed Glide SP. This does not fulfill the expectation that a deep learning model should be superior to conventional methods. This is probably because the total number of protein-ligand complexes in our training set is still too small to support training a robust deep learning model. The chemical space covered by the ligand molecules from the PDBbind data set is rather limited as compared to that covered by the molecules included in the DUD-E or LIT-PCBA data set. As a proof, the results of our UMAP analysis of those several data sets are given in Figure 6. One can see while the binders and non-binders in the DUD-E or LIT-PCBA data set are distributed sparsely over a wide space, the ligand molecules in the PDBbind general set (v.2020) tend to concentrate at certain regions. This observation is consistent with the claim by several other reports that the structural diversity of the ligand molecules in the Protein Data Bank is somewhat limited ^73, 74^. Adding the decoys selected from ChEMBL29 to our training set greatly expanded its covered chemical space to roughly the same level of DUD-E or LIT-PCBA (see Figure S2 in the *Supporting Information*). As discussed above, this treatment helps PLANET to improve its screening power (Table 4). However, we have observed that this treatment does not help PLANET much to improve its prediction of protein–ligand contact map. It is because those added decoys introduce basically all-zero labels in large number to the training set in respect to protein-ligand contact map, which makes the training set even more unbalanced for this task. What an “ideal” training set for our multi-objective model should be is of course a matter for debate. Inclusion of significantly more protein-ligand complexes with low binding affinity seems to be necessary for obtaining a more balanced training set. Such samples, however, are relatively insufficient from publicly available resources at present.

**Figure 6.**
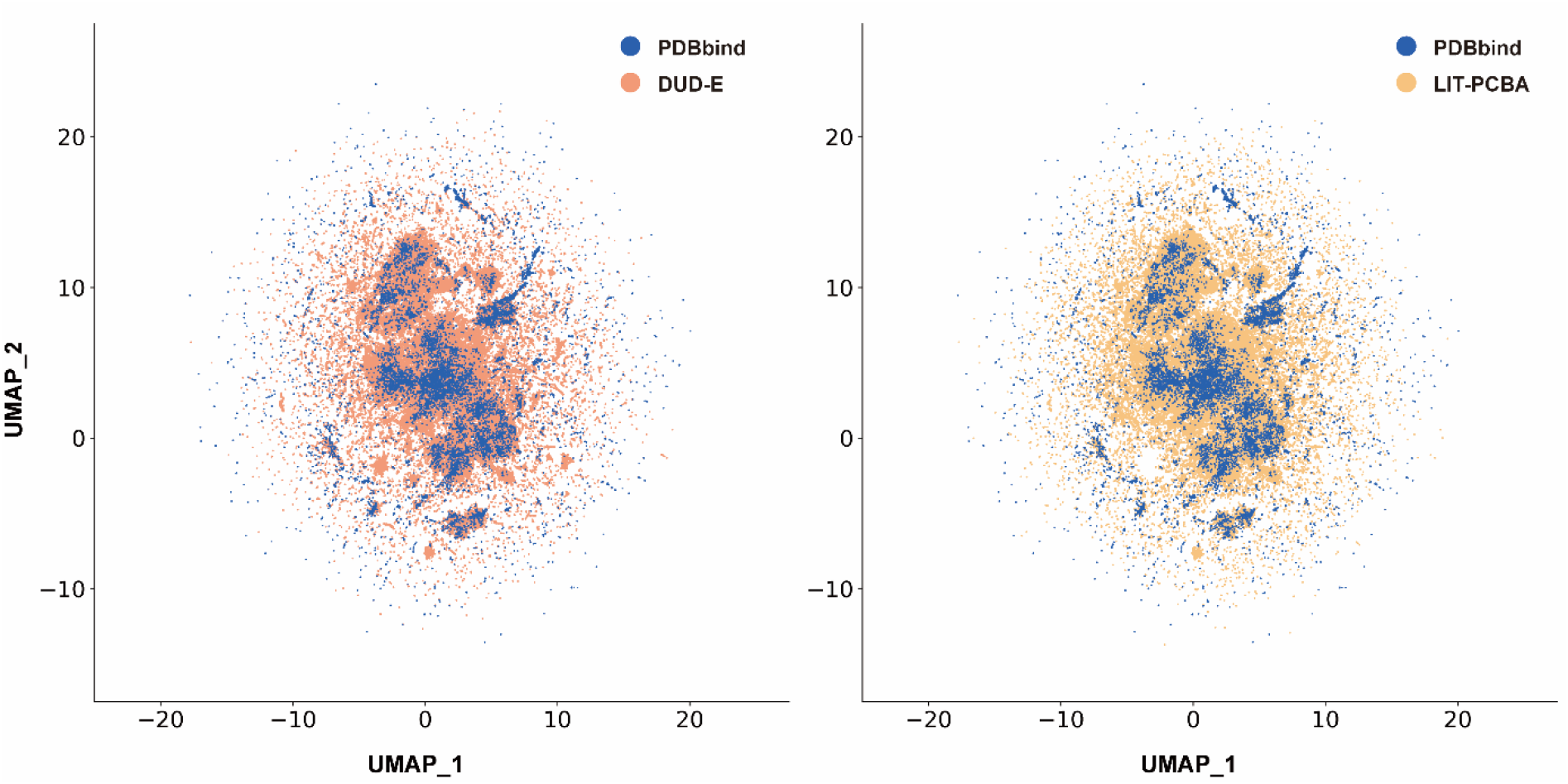
UMAP projections of the molecules in three data sets employed in our work, where blue dots represent the samples in the PDBbind general set (v.2020), salmon dots for those in DUD-E, and orange dots for those in LIT-PCBA.

We have demonstrated in this work that the multi-objective training strategy and the training set augmented with selected decoys can improve the screening power and generalizability of our PLANET model. Nevertheless, it seems that the insufficient diversity represented by the current training set sets a fundamental limit to its performance. In particular, crystal structures of protein–ligand complexes are not straightforward to obtain, so the number of such structures will not grow dramatically in the near future. From this perspective, more efficient experimental or computational methods for obtaining reliable protein–ligand complex structures are still desired. Alternatively, more capable machine learning models that can utilize small data sets in training may also provide a solution to the problem of protein-ligand binding affinity prediction.

## 4. Conclusion

In this work, we have developed a GNN-based model, called PLANET (i.e. Protein-Ligand Affinity prediction NETwork), which takes 3D structure of protein binding pocket and 2D ligand structure as input to predict protein-ligand binding affinity. Evaluation results obtained on the popular CASF-2016 benchmark indicate that the scoring power of PLANET is among the top as compared to other deep learning models. PLANET can also predict intra-ligand distance matrix and protein– ligand contact map with a fair accuracy, which makes its results more interpretable. Thus, unlike other black box–style deep learning models, our model makes prediction of protein-ligand binding affinity based on an “understanding” of protein–ligand interactions. This advantage of PLANET is further verified by the evaluation results obtained on two benchmarks designed for testing virtual screening models, i.e. DUD-E and LIT-PCBA. Here, PLANET outperforms other deep/machine learning models in terms of AUC ROC scores and EFs. Its accuracy in these virtual screening tests is generally comparable to that of the established docking method Glide SP. However, its computational speed is faster by at least two orders of magnitude because it does not need the time-consuming molecular docking process. The balanced power of PLANET should be attributed to our multi-objective training strategy and the special training set combining experimental protein-ligand complex structures and non-binder decoys. With its fair accuracy and high computational efficiency, we expect PLANET to become a practical tool for conducting large-scale virtual screening tasks in real drug discovery efforts.

## Data and Software Availability

Source codes of PLANET, as well as user manual and demo examples, are available at https://github.com/ComputArtCMCG/PLANET.

---

**Table S1.** Initial features for nodes and edges in 2D molecule graph construction

|  | Descriptors | Length |
| --- | --- | --- |
| Node (atom) | One-hot encoding for element types including C, N, O, S, P, F, Cl, Br, I, H and others | 11 |
|  | One-hot encoding for the degree of an atom including 0,1,2,3,4 and 5 or more | 6 |
|  | One-hot encoding for the formal charge of an atom including -2, -1, 0, 1 and 2 | 5 |
|  | One-hot encoding for the chiral tage of an atom including achiral R, S and unspecified | 4 |
|  | One-hot encoding for the hybridization of an atom including SP, SP2, SP3 and others | 4 |
|  | Whether the atom is aromatic or not | 1 |
| Edge (bond) | One-hot encoding for the bond type including single, double, triple and aromatic | 4 |
|  | Whether the bond is in a ring or not | 1 |
|  | Whether the bond is conjugated or not | 1 |

**Table S2.** PDB codes of protein structures used in virtual screening datasets.

| Target Name | PDB ID |
| --- | --- |
| ADRB2 | 4LDE |
| ALDH1 | 5L2N |
| VDR | 3A2J |
| ESR1_agonist | 2QZO |
| ESR1_antagonist | 5UFX |
| GBA | 2V3D |
| IDH1 | 4UMX |
| KAT2A | 5MLJ |
| MAPK1 | 4ZZN |
| MTORC1 | 4DRI |
| OPRK1 | 6B73 |
| PKM2 | 3GR4 |
| PPARG | 3B1M |
| TP53 | 3ZME |

**Table S3.** Comparison of the performance of PLANET and Glide SP on the LIT-PCBA benchmark

| Target | PLANET |  |  |  | Glide SP |  |  |  |
| --- | --- | --- | --- | --- | --- | --- | --- | --- |
|  | ROC_AUC | EF<sup>0.5%</sup> | EF<sup>1%</sup> | EF<sup>5%</sup> | ROC_AUC | EF<sup>0.5%</sup> | EF<sup>1%</sup> | EF<sup>5%</sup> |
| ADRB2 | 0.456 | 11.759 | 5.881 | 2.353 | 0.573 | 0.000 | 5.555 | 5.222 |
| ALDH1 | 0.699 | 9.085 | 11.356 | 5.000 | 0.561 | 1.933 | 2.792 | 2.320 |
| ESR1_ago | 0.575 | 15.371 | 7.685 | 4.611 | 0.513 | 13.976 | 7.106 | 2.852 |
| ESR1_ant | 0.639 | 3.882 | 6.927 | 2.593 | 0.450 | 0.234 | 0.409 | 0.631 |
| FEN1 | 0.720 | 7.596 | 6.239 | 3.913 | 0.687 | 4.367 | 4.045 | 3.384 |
| GBA | 0.707 | 4.819 | 4.819 | 2.651 | 0.581 | 5.262 | 4.209 | 2.316 |
| IDH1 | 0.522 | 0.000 | 0.000 | 2.564 | 0.430 | 0.281 | 0.562 | 0.702 |
| KAT2A | 0.513 | 4.145 | 3.109 | 1.865 | 0.490 | 1.005 | 1.507 | 1.106 |
| MAPK1 | 0.662 | 0.651 | 1.301 | 2.280 | 0.558 | 1.336 | 1.289 | 1.429 |
| MTORC1 | 0.443 | 0.000 | 0.000 | 0.618 | 0.490 | 0.842 | 0.842 | 1.106 |
| OPRK1 | 0.398 | 0.000 | 0.000 | 1.667 | 0.528 | 0.000 | 4.166 | 1.667 |
| PKM2 | 0.596 | 1.830 | 1.465 | 1.135 | 0.669 | 5.942 | 4.020 | 2.762 |
| PPARG | 0.728 | 0.000 | 3.660 | 2.961 | 0.670 | 4.064 | 3.879 | 4.436 |
| TP53 | 0.541 | 9.772 | 5.000 | 1.514 | 0.640 | 4.635 | 2.369 | 1.921 |
| VDR | 0.398 | 0.682 | 0.569 | 0.705 | 0.462 | 0.217 | 0.433 | 0.520 |

**Figure S1.**
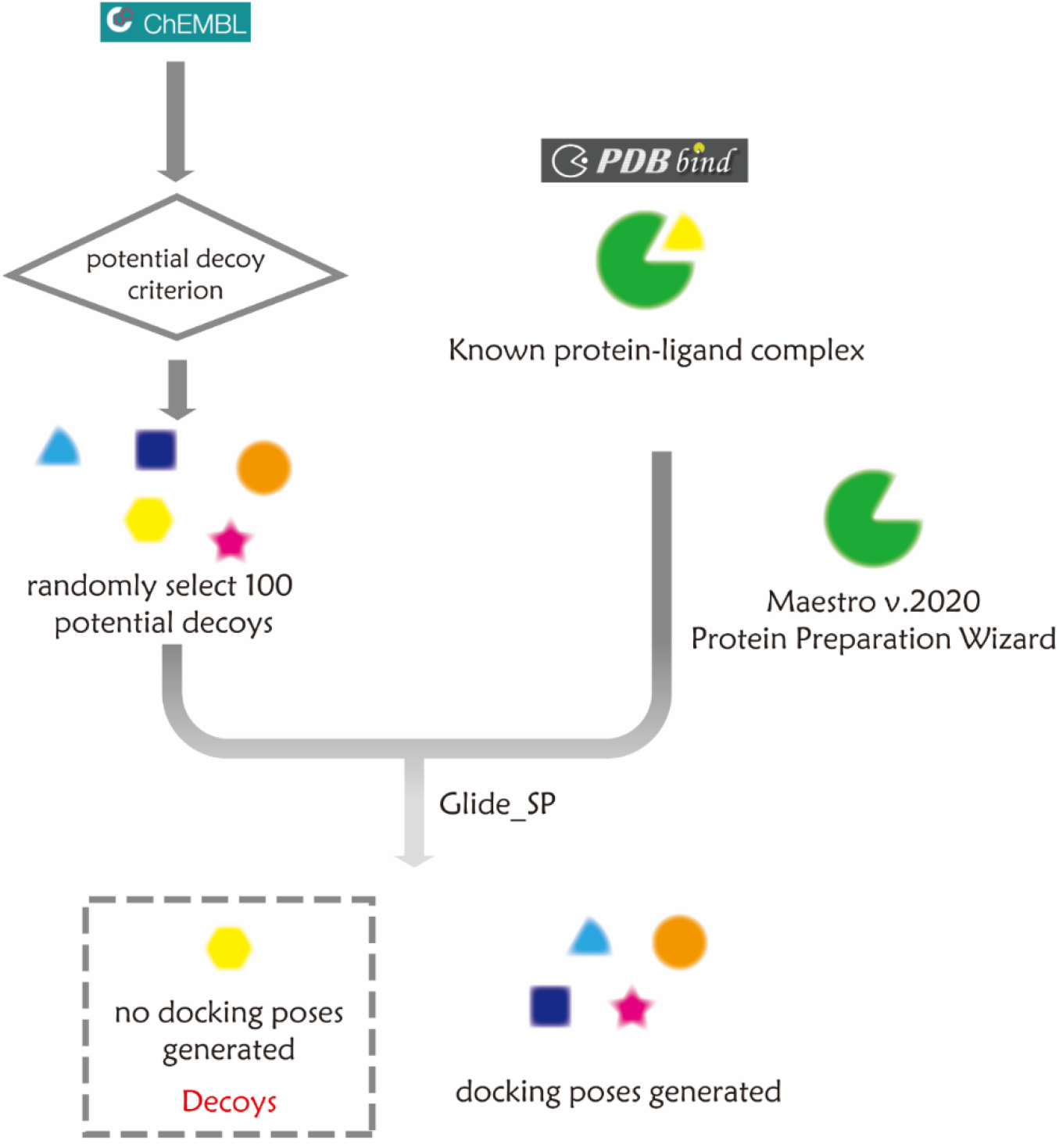
Workflow of generating decoys for augmenting dataset. Molecular properties were calculated for all 2.1 million molecules in ChEMBL 29. For each protein-ligand complex in our training set, candidate decoys were required to meet at least six of the following eight criteria: (1) Difference in molecule weight was less than 10%; (2) Difference in log*P* is less than 1 units; (3) Difference in number of hydrogen bond donor was less than 1; (4) Difference in number of hydrogen bond acceptor was less than 1; (5) Difference in number of chiral centers was less than 1; (6) Difference in number of rings was less than 1; (7) Difference in number of heavy atoms was less than 1; (8) Difference in number of total atoms was less than 1. Note that the qualified candidates were further examined by docking them into the same binding site as the ligand molecule in the given protein-ligand complex. The candidates for which Glide SP failed to generate any valid binding pose were finally accepted as the non-binder decoys for the given protein-ligand complex.

**Figure S2.**
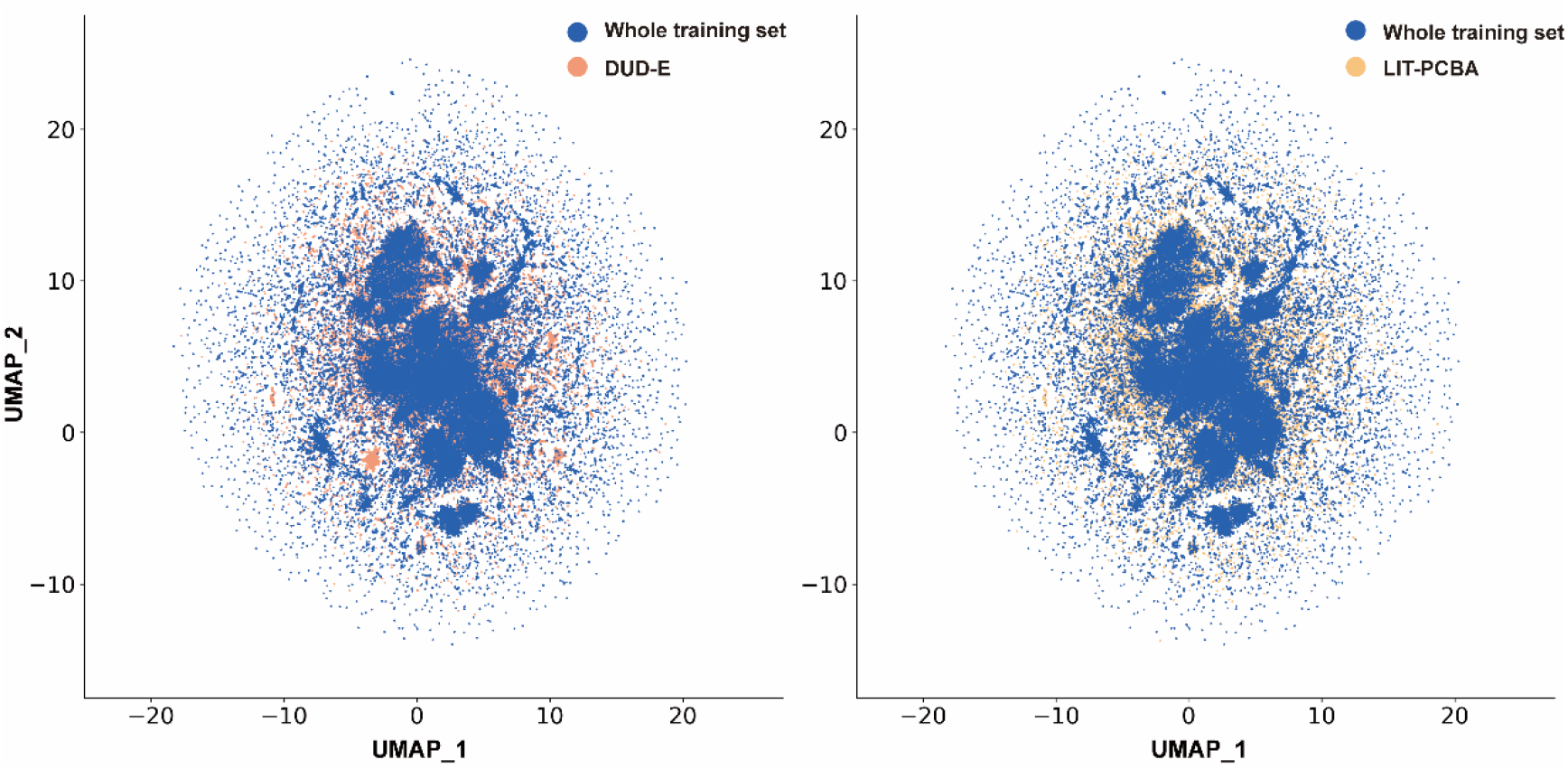
UMAP projection of all molecules in our training set, including the ligands in the PDBbind general set (v.2020) and those selected decoys from ChEMBL 29.

